# Distributional changes in myelin-specific MRI markers uncover dynamics in the fornix following spatial navigation training

**DOI:** 10.1101/2020.12.13.422557

**Authors:** Debbie Anaby, Benjamin C. Tendler, Matthias S. Treder, Simon Hametner, Ulrike Koeck, Chantal M.W. Tax, Greg D. Parker, Daniel Barazany, Yaniv Assaf, Derek K. Jones

**Affiliations:** Cardiff University Brain Research Imaging Centre (CUBRIC), Cardiff University, Cardiff CF10 3AT, United Kingdom; Department of Diagnostic Imaging, Sheba Medical Center, Tel HaShomer 52621, Ramat Gan, Israel; Wellcome Centre for Integrative Neuroimaging, FMRIB, Nuffield Department of Clinical Neurosciences, University of Oxford, Oxford OX3 9DU, United Kingdom; Sir Peter Mansfield Imaging Centre, School of Physics and Astronomy, University of Nottingham, United Kingdom; School of Computer Science & Informatics, Cardiff University, Cardiff CF24 3AA, United Kingdom; Division of Neuropathology and Neurochemistry, Department of Neurology, Medical University of Vienna, Austria; Neuroimmunology Department, Center of Brain Research, Medical University of Vienna, Vienna 1090, Austria; Experimental MRI Centre (EMRIC), School of Biosciences, Cardiff University, Cardiff CF10 3AX, United Kingdom; Strauss computational neuroimaging center, Tel Aviv University, Tel Aviv 69978, Israel; Department of Neurobiology, George S. Wise Faculty of Life Sciences, Tel Aviv University, Tel Aviv 69978, Israel; Sagol School of Neurosciences, Tel Aviv University, Tel Aviv 69978, Israel; Mary McKillop Institute for Health Research, Faculty of Health Sciences, Australian Catholic University, Melbourne, Victoria 3065, Australia

## Abstract

Increasing evidence implicates white matter (WM) dynamics supporting learning in the mature brain. Recent MRI studies, mostly using diffusion tensor MRI (DT-MRI), have demonstrated learning-induced WM changes at the microstructural level. However, while DT-MRI-derived measures have sensitivity to general WM microstructural changes, they lack compartmental specificity, making them difficult to relate to underlying cellular mechanisms, stymying deeper understanding of mechanisms supporting training-induced gains in performance. Gaining a deeper understanding demands a more detailed characterization of changes in specific WM sub-components. To this end, four microstructural MRI techniques were employed to study alterations in rat brains after 5-days of water maze training: DT-MRI; Composite Hindered and Restricted Model of Diffusion (CHARMED); magnetization transfer (MT) imaging; quantitative susceptibility mapping and R_2_*.

The hypothesis tested here was that microstructural changes would be: (i) observed in tracts supporting spatial navigation, i.e., fornix and corpus callosum (CC); and (ii) more pronounced in the myelin-specific measures.

Medians and distributions of microstructural parameters were derived along the fornix, CC and cingulum (as a comparison tract) using the ‘tractometry’ approach. Summary measures were derived from different metrics using unsupervised data reduction. Significant pre-*vs*-post training differences were found in the medians of two principal components loading on: (i) anisotropy indices; and (ii) MT ratio. The most striking effect, however, was seen in the *distributions* of pre-*vs*-post training MT ratio in the fornix, consistent with the primary hypothesis, and highlighting the value of this alternative to the standard approach (i.e., comparing means/medians of DT-MRI parameters) for studying neuroplasticity *in vivo*.

**Significance statement:** Recent MRI-studies have demonstrated that white matter (WM) dynamics support learning, even in the mature brain. However, most studies are based on diffusion tensor MRI (DT-MRI) measures, which although providing sensitivity to WM microstructural changes, do not provide a direct translation to the underlying cellular mechanisms making interpretation difficult. Using alternative quantitative MRI approaches, this study provided more specific subcomponent microstructural insights into the brain’s WM response to water-maze training. We show that myelin-specific MR measures show more marked changes than axonal-specific and DT-MRI measures, in WM tracts responsible for spatial navigation. The results promote adoption of these alternative approaches to DT-MRI for studying neuroplasticity *in vivo*.

## Introduction

It has been recently demonstrated, using MRI, that white matter (WM) in the mature brain is dynamic, and that these dynamics contribute to learning (Johansen-Berg et al., 2007; Blumenfeld-Katzir et al., 2011; Hofstetter et al., 2013; Caeyenberghs et al., 2016; Hofstetter and Assaf, 2017; Metzler-Baddeley et al., 2017). Numerous mechanisms supporting learning have been demonstrated at the cellular level. Sampaio-Baptista et al. (2013) showed increased myelin-basic protein (MBP) immunolabeling in rats after training, suggesting that learning triggers the formation of new myelin, while McKenzie et al. (2014) demonstrated that generation of new oligodendrocytes (OLs) is crucial for learning motor skills.

Most neuroplasticity MRI studies use diffusion tensor MRI (DT-MRI), reporting increased fractional anisotropy (FA) and reduced mean diffusivity (MD) in WM after long-term training. For example, Blumenfeld-Katzir et al. (2011) found increased FA in the rat corpus callosum after spatial learning in a water maze. Others, however, found the opposite effect, with lower FA / higher MD after training (Hänggi et al., 2010; Taubert et al., 2010). While differences in training tasks might explain these differences in imaging findings, it is also well-known that DT-MRI-derived measures are degenerate and therefore difficult to interpret. In addition to myelin, (Beaulieu, 2002), FA is modulated by fiber architecture (Basser and Pierpaoli, 1996; Douaud et al., 2011), orientation dispersion (Budde and Annese, 2013), axon density and axon diameter distribution (Takahashi et al., 2002; Shemesh, 2018). Thus, FA changes are not specific to myelin (or any other sub-component). Moreover, Tyszka et al.’s (2006) study on Shiverer mice, a model characterized by almost complete absence of compact myelin in the central nervous system (Readhead and Hood, 1990), showed that FA was only 15% lower compared to wild-type mice, indicating FA’s low sensitivity to subtle differences in myelin.

Given emergent evidence that myelin changes are critical for learning, and hypothesized as the driving factor underpinning previously reported learning-induced FA changes, an individual characterization of myelin changes *in vivo* should be considered alongside other WM sub-components. Such a characterization requires alternative MRI acquisition and modelling approaches. For example, quantitative magnetization transfer (qMT) is associated with the relative amount of myelin in WM (Turati et al., 2015) susceptibility weighted imaging (SWI) provides complementary information on components with differential magnetic susceptibility including myelin and iron (Haacke et al., 2015), while the Composite Hindered and Restricted Model of Diffusion (CHARMED) (Assaf et al., 2004; Assaf and Basser, 2005) provides more compartment-specific diffusion (e.g., intra-axonal, extra-axonal) parameters. Here we combine four complementary MRI techniques, i.e., DT-MRI, CHARMED, magnetization transfer (MT) and SWI via R_2_*- & quantitative susceptibility mapping (QSM) to characterize specific WM alterations in the rat brain after spatial working-memory training. Histological assessment of myelin was also performed on a sample of pre- and post-training rat brains.

Our hypotheses were that microstructural WM changes would be:

i. found in the fornix, the major output tract from the hippocampus, which is critical for spatial navigation (Hodgetts et al., 2019). We also investigated the corpus callosum, given evidence of post-training changes (Blumenfeld-Katzir et al., 2011) and the cingulum, which was not expected to change and therefore serves as an internal comparison tract; and
ii. more marked in the myelin-specific measures (e.g., MT) than the axon-specific measures (e.g., CHARMED), based on previous evidence that myelin is crucial in training-induced WM changes.

Note: although our hypotheses are focused on myelin, we have included measures that are more sensitive to differences in axonal characteristics. This is because most *in vivo* microstructural plasticity studies to date have used DT-MRI, which cannot distinguish between axon or myelin changes and is sensitive, to a lesser or greater extent, to both. Our aim here was to break down the plasticity response to microstructural subcomponents and demonstrate *specificity* of changes to one compartment. Demonstrating specificity requires isolation of the compartment that we <u>do</u> expect to change, and the compartment that we do not expect to change.

## Materials and Methods

### Ethics statement

This study was approved by the Tel Aviv University Committee on Animal Care and Use and conducted according to the guidelines for research involving animals (permit number: L-04-16-009).

### Experimental design

18 male Wistar rats, 2.5 months of age were studied. All rats were maintained on a 12-h light/12-h dark cycle with access to food and water *ad libitum*. 12 rats underwent two MRI scans, before and after 5 days of learning and memory training in a Morris water maze (Duval et al., 2018). Prior to scanning, the rats were anesthetized with 1-2% isoflurane in oxygen. 3 rats underwent the first scan and were then perfused for histology staining. Morris water maze training was performed in a 120 cm diameter pool in which a platform was hidden in one of four quadrants. Each rat underwent two rounds of training per day, with a 30 minutes rest period between the two rounds. Each training round comprised four swims, in which the rats were placed at different quadrants of the pool and given 1 minute to find the platform. Rats that did not succeed in finding the platform within this time limit were led to it. All rats were given 10 seconds to stand on top of the platform and then taken out of the pool to rest.

### Imaging

MRI was performed with a 7T/30 Bruker MRI scanner (Bruker, Karlsruhe, Germany) equipped with 400 mT/m gradients. A body coil (outer/inner diameter of 112/72 mm) was used for excitation and a quadrature coil (15 mm diameter) was used as a receiver.

The scanning protocol comprised three different types of acquisition:

i. **2D diffusion-weighted pulsed-gradient spin-echo (DW-PGSE) – for diffusion MRI:** isotropic image resolution of 370 μm (32 axial slices), TR/TE= (8s/28ms), NEX = 1, b-values = 1000, 2000 and 4000 s/mm^2^, Δ/δ = (14 ms/7.5 ms), 30 noncollinear directions;
ii. **3D multi-gradient echo (MGE) – for susceptibility-sensitive imaging:** image resolution of 185×185×370 μm, 8 TEs (first TE=3.4 ms, echo-spacing=5.6 ms);
iii. **3D fast low angle shot (FLASH) – for magnetization transfer imaging:** FLASH with magnetization transfer saturating Gaussian pulses, 2 flip angles of 1000° and 2800° with 12 offsets (ranging between 1000 and 30000 Hz), 3 images with no saturating pulses using isotropic resolution of 370 μm. Note that full quantitative MT measures were not calculated due to a corruption of the quantitative T1 maps (needed for the full quantitative MT model), therefore, the magnetization transfer ratio (MTR) was calculated using one of the frequency-offsets collected with the qMT protocol.

The duration of the complete acquisition protocol was 1.5 hours.

### MRI image analysis

Each MR contrast required a dedicated processing pipeline using custom MATLAB (The Mathworks) scripts:

i. **DW-PGSE data** The diffusion-weighted data were analysed in multiple ways.

a. <u>DT-MRI Analysis</u>: Using the data collected with the first shell (b=1000 s/mm^2^), the diffusion tensor was estimated robustly in each voxel using the RESTORE algorithm (Chang et al., 2005) in the *ExploreDTI* software package (Leemans et al., 2009). From the tensor, four scalar measures were derived, namely: fractional anisotropy (FA), mean diffusivity (MD), radial diffusivity (RD) and longitudinal diffusivity λ_1_.
b. <u>CHARMED analysis</u>: The CHARMED model (Assaf et al., 2004; Assaf and Basser, 2005) was fit to all shells of the diffusion-weighted data. This model assumes that two water populations are present; intra-axonal water that exhibits ‘restricted’ diffusion and extra-axonal water that exhibits ‘hindered’ diffusion. The parameter of interest here was the ‘restricted diffusion signal fraction’ (Fr), i.e. the fraction of the signal that arises from the restricted diffusion population. (Note: Fr is often referred to as ‘axon density’).
c. <u>Tractography analysis</u>: The RESDORE algorithm (Parker et al., 2013) was applied on the b=2000 s/mm^2^ shell data to obtain robust estimates of the fibre orientation density function (fODF) using the damped Richardson-Lucy (dRL) algorithm (Dell’Acqua et al., 2010). These fODFs served as input to deterministic tractography to reconstruct the fornix, CC and cingulum tracts (bilaterally), followed by manual delineation of ‘way-points’ in each data set by an operator blinded to ‘pre- vs. post-training status’ to avoid bias (see Figure S1 in Supplementary Information). The fornix and CC were hypothesized *a priori* to change markedly due to training, whereas the cingulum was not, and is therefore referred to here as a comparison tract. (N.B. We do not refer to the cingulum as a ‘control’ tract because all parts of the brain were subjected to the same learning procedure).
ii. **MGE Data** Susceptibility and R_2_* maps were derived from the MGE data. R_2_* maps were generated from the gradient echo magnitude data assuming mono-exponential signal evolution. For QSM, frequency maps were first estimated from the gradient echo phase by fitting to the complex signal across all echoes (Liu et al., 2013), followed by background field removal using v-SHARP (Wu et al., 2012). Quantitative susceptibility maps were subsequently generated following a similar approach to (Rochefort et al., 2010), using a magnitude-weighted least squares minimisation (regularised using both the brain mask and susceptibility gradients estimated via fastQSM (Li et al., 2015). The susceptibility in each animal, at each time point, was referenced to the average of a CSF chosen region.
iii. **FLASH Data** The magnetization transfer ratio (MTR) was computed according to the formula: MTR = (S_MT_ – S_0_)/S_0_, where S_0_ is the signal intensity without any off-resonance pulse applied, and S_MT_ is the signal intensity obtained with an off-resonance pulse with offset of 1860 Hz and 1000° flip angle.

### Tractometry

Maps of each of the derived parameters: fractional anisotropy (FA), mean diffusivity (MD), radial diffusivity (RD), longitudinal diffusivity λ_1_, restricted diffusion signal fraction (Fr), MTR, R_2_* and susceptibility were co-registered within each rat (using SPM8, version 6313 (Penny et al., 2007)) and then projected on to the reconstructed tract-bundles, following the ‘Tractometry’ approach (Bells et al. 2011). For each parameter, the median value of all vertex-wise estimates was computed and the distribution stored for later analyses. De Santis et al., 2014, previously deployed tractometry in healthy human subjects, and uncovered covariances between the different parameters. This motivated a principal component analysis (PCA) in the present study, to capture the principal sources of variance. This approach was recently shown to improve the disentangling of neurobiological underpinnings of WM organization (Chamberland et al., 2019). PCA was performed as follows: to bring pre- and post-training measures on equal footing, they were first normalized by their mean and standard deviation (averaged across the three different anatomical bundles). Covariance matrices were then calculated for each white matter pathway, normalized by the trace of the covariance matrix, and pooled prior to principal component decomposition. Principal components (PCs) were calculated using MATLAB’s *pcacov* function. The subset of PCs explaining at least 95% of the total variance was selected. Varimax rotation was then performed to improve the interpretability of the result.

### Power Analysis

The sample size needed to obtain a statistical power of 0.8 with an effect size of 5% was calculated to be 11 rats per group, based on a 1-way analysis of variance (ANOVA) pairwise, 2-sided equality statistical model (Shein-Chung et al., 2008).

### Statistical Analysis

Following the strict recommendation of Thomas and Baker (Thomas and Baker, 2013) for making robust statistical inferences in MRI-based training studies, a significant interaction should be demonstrated, while any other specific effect of training provides only weak evidence of training-induced changes. Consequently, all PCs were subjected to a 2-way repeated measures analysis of variance (ANOVA) for main effects of ‘Tract’ and ‘Time’ and their interaction (SPSS statistics software), to provide evidence of anatomical specificity of results. Only in the event of a significant Tract × Time interaction were *post-hoc* paired t-tests performed to look for significant effects of training on microstructural parameters. As a final step in the analysis, the pre-training vs post-training *distributions* of the derived measures (i.e., taking all the vertex-wise measures of streamlines for the whole reconstructed tract), rather than the median values, were compared using a two-sample Kolmogorov-Smirnov test and significance testing performed using permutation testing (1000 permutations of pre-vs-post group membership), both using in-house MATLAB scripts.

### Histology and histological analysis

For the histology analysis, a total of 6 rats (3 pre-training and 3 post-training) were perfused with PBS (1M) and 2.5% Glutaraldehyde and then placed in a vial of 2.5% Glutaraldehyde diluted in PBS. After fixation and embedding in paraffin, blocks were cut with a microtome and 3-4 μm thick sections were mounted on glass slides. For two rat brains (one pre-training, one post-training), 5 steps with 50-micron gaps through the investigated anatomical structures were derived. One section per step was stained for these animals. For the other 4 rat brains, 5 steps with 50-micron gaps and 3 directly consecutive sections per step were made. All sections were stained for Luxol-fast blue-periodic acid Schiff (LFB-PAS). The identical staining protocol has been previously applied for the comparison with quantitative susceptibility mapping (Hametner et al., 2018) and myelin water imaging (Birkl et al., 2019). Sections were digitized with a Hamamatsu Nanozoomer slide scanner at 200x magnification. Images were colour-deconvolved using the colour deconvolution plugin for ImageJ (Ruifrok and Johnston, 2001), applying the vector “H-PAS”. The resulting blue-channel and red channel images were converted to an 8-bit grey scale image and inverted. The blue-channel image represents the Luxol fast blue dye for myelin, while the red-channel image depicts the PAS counterstain, which yields a pink staining of the neocortical neuropil. Blue-channel image intensities were taken for the myelin density estimates. Red-channel image intensities were used as surrogate for the slice thickness for normalization of myelin estimates, based on the observation and assumption that neocortical PAS staining in the studied animals is mainly a variable of slice thickness. Red channel intensities for normalization were invariably taken from small regions of the temporal neocortex. Myelin intensities were taken from the corpus callosum, the white matter directly subjacent to the cingular cortex, and the fimbria hippocampi (shown in Figure 1). All intensities were normalized for each section with the respective red channel intensities of the same slice. Normalized intensities were averaged for all slices such that one data point was retrieved per region per rat. We then performed a 2-tailed t-test for significance.

**Figure 1:**
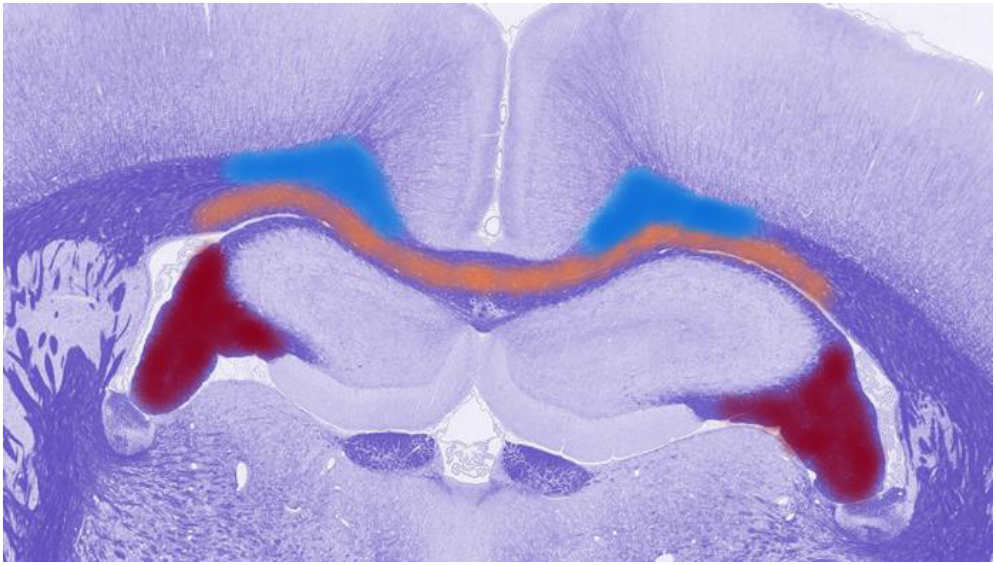
Three WM regions that were considered for histology analysis; subjacent to the fimbria hippocampi (‘fornix’) (red), corpus callosum (orange) and subjacent to the cingular cortex (‘cingulum’) (blue).

## Results

Figure 2 shows the ‘time to platform’ in each swim. The average time to reach the platform in the first day (2 rounds) was ~33 sec with marked reductions throughout the following days reaching ~7 sec on the fifth day, demonstrating that learning was occurring.

**Figure 2:**
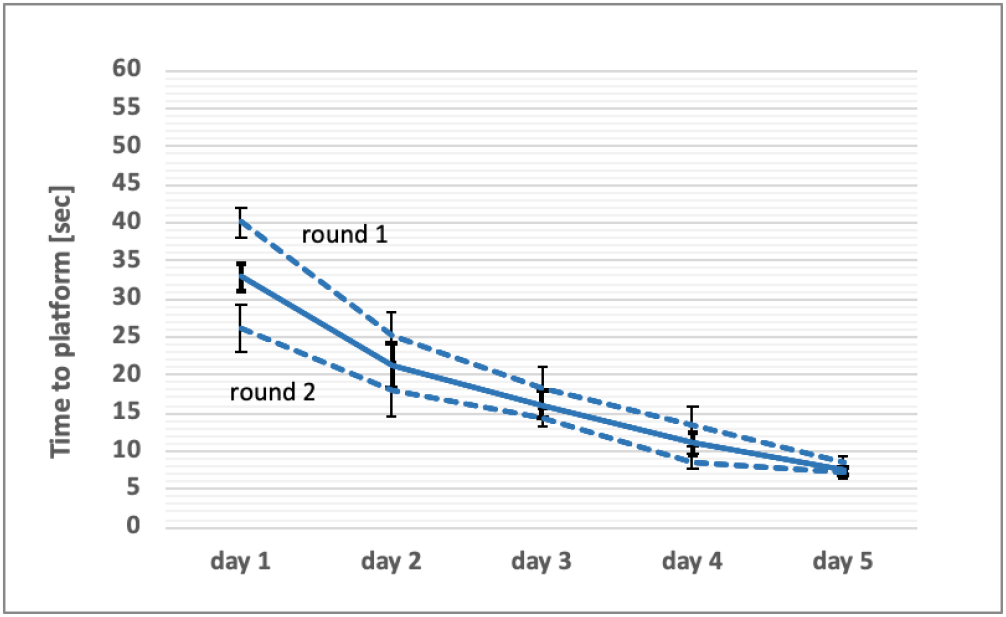
Mean ‘time to platform’ in rounds 1 and 2 (dotted lines) and their average (solid line) for each of the 5 training days.

Eight microstructural measures were derived from the scanning protocol; FA, MD, RD, λ_1_, Fr, MTR, R_2_* and quantitative susceptibility, χ. Figure 3 shows cross-correlation matrices in the three tracts along with specific correlations of three pairs of measures in the fornix, CC and cingulum. Several pairs of measures showed significant covariance, motivating a principal component analysis (PCA).

**Figure 3:**
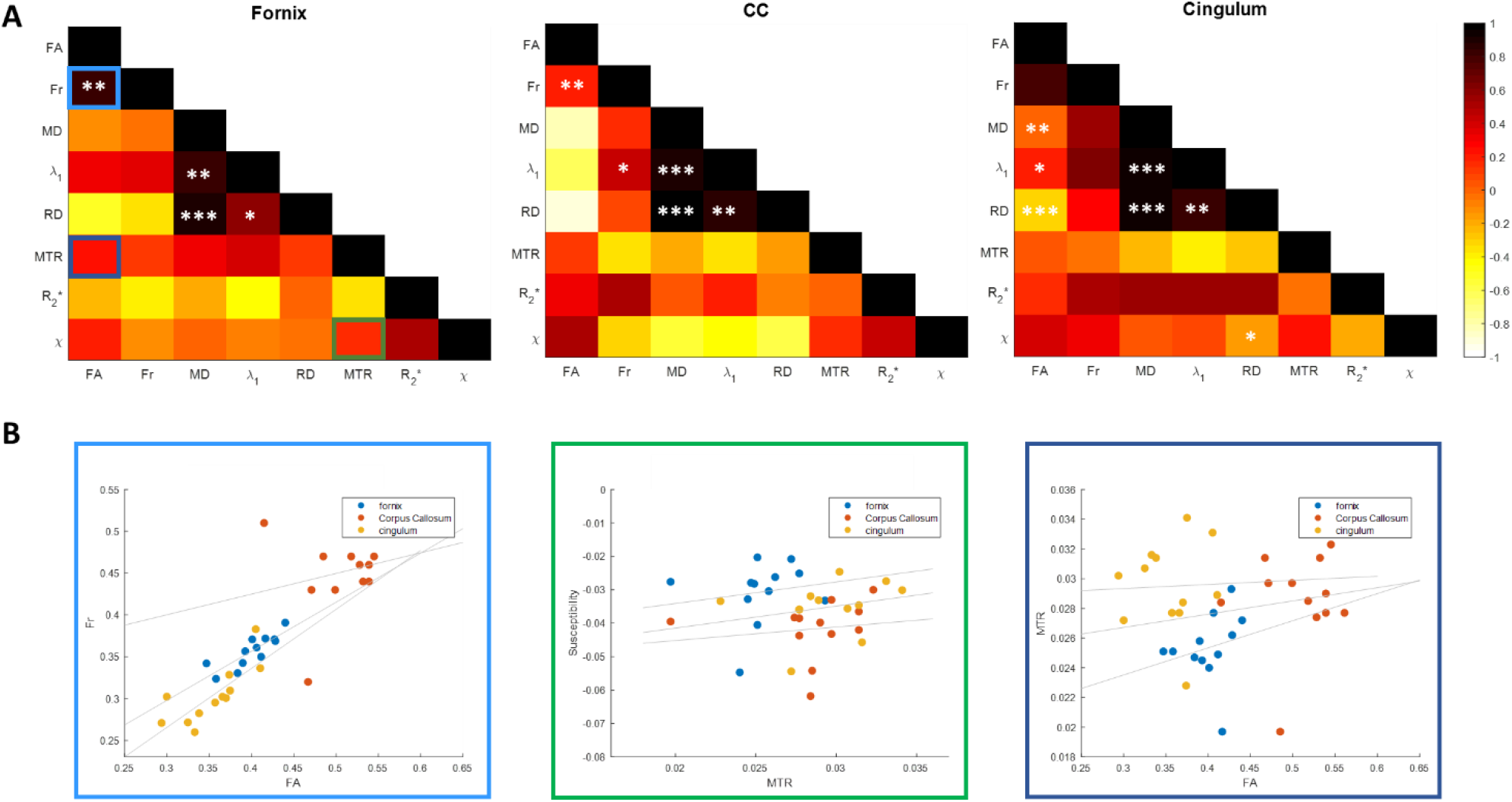
(A) Cross correlation matrices of the eight measures in fornix, CC and cingulum. Significant pairs of measures are marked by *p<0.05, **p<0.005, ***p<0.0005 (B) Correlations of FA vs. Fr; fornix - R=0.84, p=0.0006 and cingulum - R=0.77, p=0.0032 (left plot), MTR vs. Susceptibility; no significance (middle plot) and FA vs. MTR; no significance (right plot) in fornix, CC and cingulum.

The PCA yielded 5 principal components (Figure 4) that explained over 95% of the variance; The first and second components were dominated by diffusion measures (explaining variance of ~34% and 22%, respectively). The third, fourth and fifth PCs are each dominated by a single measure: susceptibility (explained variance ~16%); MTR (explained variance ~14%); and R_2_* (explained variance ~12%), respectively.

**Figure 4:**
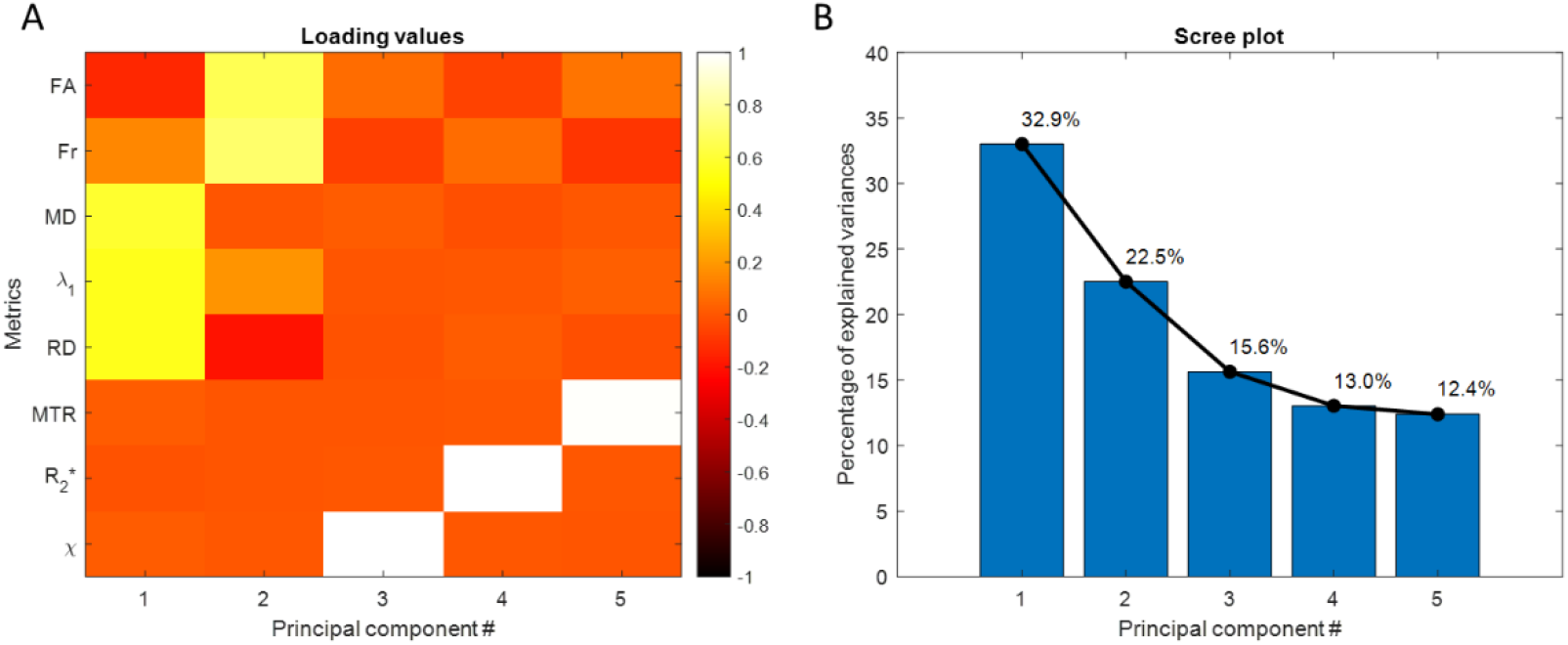
(A) Absolute loading values of the eight microstructural indices; rows represent the absolute loading values of the different measures and columns represent the principal components. (B) Scree plot representing the percent of explained variances.

Figure 5 shows the average PC scores for each tract at the two time points, along with the loading values for each PC. Results from the 2-way ANOVA statistical tests showed that the main effect of ‘Tract’ was significant for all PCs. Taking each PC in order of percentage variance explained:

**Figure 5:**
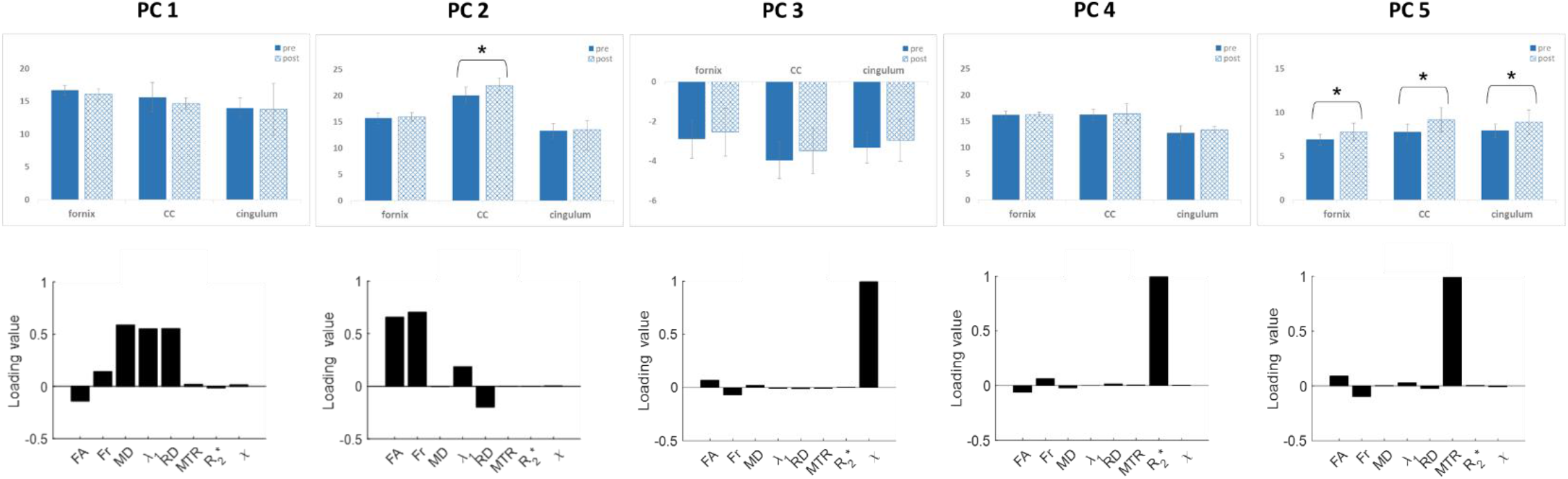
Average PC scores at the two time points of each tract along with the loading values for each PC. Bonferroni-Holm corrected p-values derived from paired t-tests between the two time points within each tract in every PC.

PC_1_, which is dominated mainly by diffusivity measures (MD, RD and λ_1_), showed a non-significant reduction between the pre- and post-training time points in fornix and corpus callosum (this PC was relatively stable between the two time points in the cingulum). No statistical significance was found other than the main effect of ‘tract’.

PC_2_, which is dominated mainly by anisotropy measures (FA and Fr), showed an increase between pre- and post-training median values in the CC, while values in the fornix and cingulum were relatively constant. There was a significant ‘Tract × Time’ interaction (F(2,22)=3.54, p=0.047, η_p_^2^=0.243), and main effect of ‘Time’ (F(1,11)=9.74, p=0.01, η_p_^2^=0.47). Post-hoc paired t-tests showed a significant effect of training only in the CC (p=0.007). These results effectively replicate the result of Blumenfeld-Katzir et al. 2011 who also showed a significant increase in FA (derived from DT-MRI) in the CC of rats in a similar water maze study.

PC_3_, which loaded almost exclusively on susceptibility, shows increasing trends between the pre- and post-training values for all three tracts. However, no statistical significance was found other than for main effect of ‘Tract’. The same was true for PC_4_, dominated by R_2_*.

PC_5_, which is dominated by MTR, showed a significant ‘Tract × Time’ interaction (F(2,22)=5.53, p=0.01, η_p_^2^=0.335), a main effect of ‘tract’ (F(2,22)=79.8, p=0.0001, η_p_^2^=0.879), and a main effect of ‘Time’ (F(1,11)=5.4, p=0.04, η_p_^2^=0.33). The *post-hoc* paired t-tests in each of the three tracts showed a significant effect of training (after Bonferonni-Holm multiple correction) in all three tracts: fornix (p=0.019), CC (p=0.009) and cingulum (p=0.049). Note, however, that the significance of change in the cingulum is borderline.

Figure 6 shows Kolmogorov-Smirnov test (K-S) statistic values (maximum distance between the two cumulative density functions) comparing pre- and post-training *distributions* in 5 microstructural measures for every rat. Its values for the MTR in the fornix are markedly higher than for all other measures in all tracts, indicating quantitatively that this myelin-sensitive marker shows the greatest shift in distribution between pre- and post-training state in the fornix.

**Figure 6:**
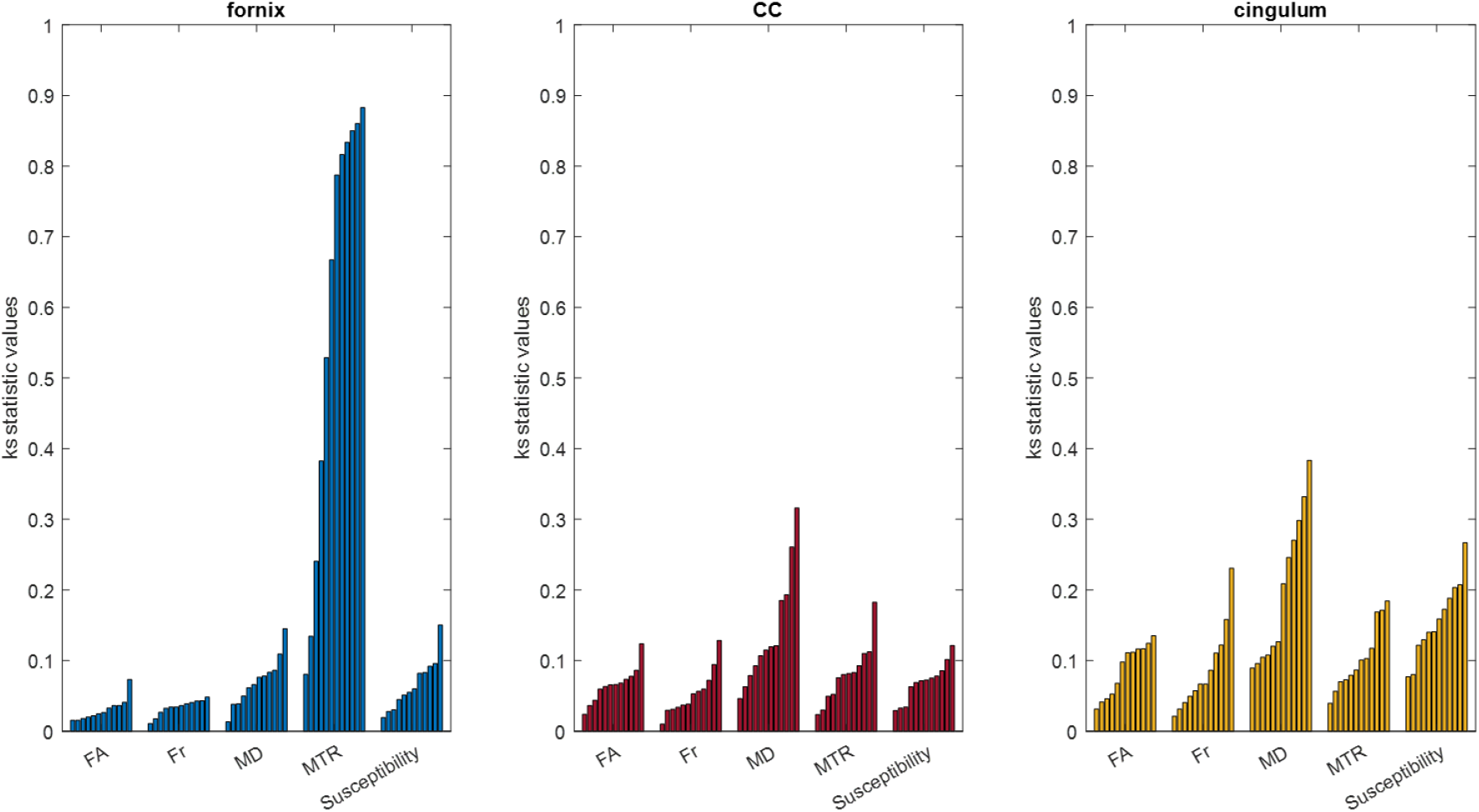
Kolmogorov-Smirnov test-statistic values of pre- vs. post-training distributions of 5 measures; FA, Fr, MD, MTR and Susceptibility, for every rat, in the fornix, CC and cingulum.

Moreover, there are clearly large inter-individual differences in the way that a rat’s microstructure responds to training. Permutation testing revealed that the K-S statistic was significant for all measures in all tracts at p=0.05 (uncorrected). This high significance most likely arises due to the large number of data points being compared in the distribution.

To complement the K-S statistics shown in Figure 6, Supplementary Figure S2 shows individual changes (pre- vs post training) in median MTR values in each rat.

Figure 7 shows the average normalized intensities of the myelin-stained image, in the three WM regions of interest in the pre and post-training groups (see Table S1 in Supplementary Material for individual values). It is important to note that each group comprises three different rats, and the histological analysis is, for obvious reasons, not longitudinal. Moreover, individual differences in myelin-stained image intensity at baseline (pre-trained rats) were present (coefficient of variation = 5%), further challenging any pre- vs post-training inferences. Finally, the number of animals was low (3 in each group). Despite these limitations, and although not significant using a 2-tailed t-test, we found higher values on average of the myelin stain in the post training group compared with the pre-training group, in all three WM regions.

**Figure 7:**
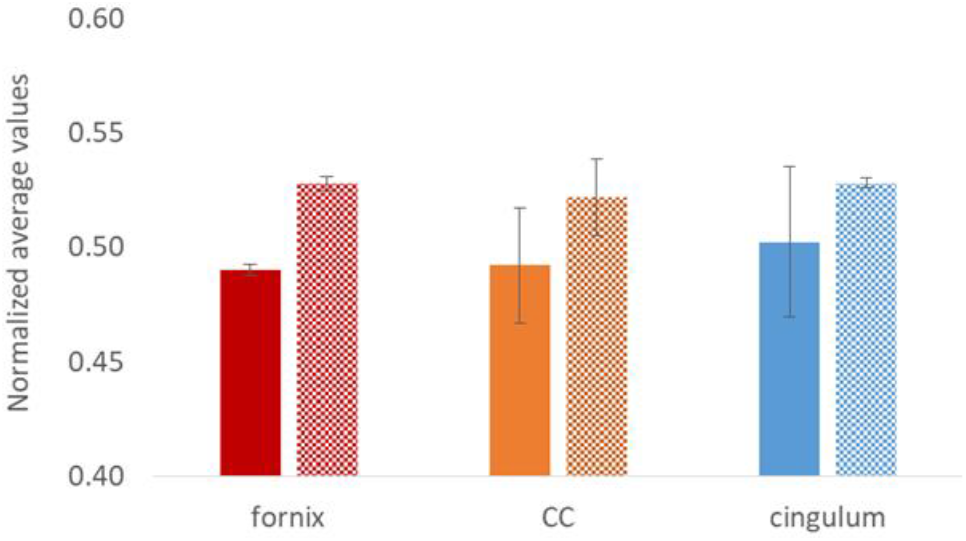
Normalized average values of histology myelin stain in pre- (solid filled) and post-training (texture filled) groups.

## Discussion

The brain’s microstructural response to learning or enriched environments has been reported previously. Changes in synapse and astrocyte number and morphology have been demonstrated (Markham and Greenough, 2004; Stuchlik, 2014). Oligodendrocytes (OLs) and myelination changes have also been reported in several studies, with Szeligo and Leblond, (Szeligo and Leblond, 1977), being the first to report increased OL count in the visual cortex of rats raised in an enriched environment. This observation was repeated by Sirevaag and Greenough (Sirevaag and Greenough, 1987), and was followed by two studies reporting: (i) an increase in the number of myelinated axons in the rat splenial CC (Juraska and Kopcik, 1988); and (ii) an increase in the size of monkey CC (Sánchez et al., 1998), in response to an enriched environment. Further, an important study in mice by McKenzie et al. showed that learning a new motor skill induced production of newly formed OLs. Critically, blocking production of these new OLs during adulthood, while maintaining pre-existing OLs and myelin, prevented the mice from mastering the task. Of high relevance to the current study, this was taken as direct evidence that new OLs, and therefore <u>new myelin</u>, are essential to learning new motor skills (McKenzie et al., 2014).

In this study we use advanced MRI methods to complement DT-MRI with the aim of providing more specific information regarding changes in WM microstructural subcomponents that may occur in rat brains after water maze training. Our main hypothesis was that myelin-specific MR measures would show more marked changes than other measures (DT-MRI and ‘axonal-specific’ measures), and that this would be most evident in tracts supporting spatial navigation, predominantly in the fornix (Hofstetter et al., 2013; Hofstetter and Assaf, 2017) and, following (Blumenfeld-Katzir et al., 2011) the CC.

Indeed, in addition to changes in the fornix, we did see changes in the CC as demonstrated by both MTR and anisotropy indices (mainly FA and Fr). As noted above, increased FA has also been reported in the CC of rats after water maze training (Blumenfeld-Katzir et al., 2011). CC changes after working memory training have also been reported. For example, Takeuchi et al. (2010) showed a significant change in WM regions adjacent to the dorsolateral prefrontal cortex (DLPFC), the anterior part of the body of the CC and the genu of the CC, after a working memory training. The bilateral DLPFCs are the key nodes of working memory such that enhanced WM microstructure in those CC regions may suggest increased interhemispheric information transfer between them.

In the fornix itself, we observed significant training-induced changes solely in the MTR. This was apparent when comparing just the median of the values along a given tract. However, Figure 6 shows that collapsing the data in this way neglects a wealth of information about the *distribution* of values along the tract. Indeed, when comparing the distributions between pre- and post-training, we see striking differences in the pre- vs post-training distributions in the MTR in the fornix, entirely consistent with our hypothesis. This highlights the importance of not simply relying on summary statistics (mean/median) as is done in most MRI studies of white matter plasticity, for analyzing training-induced changes, and the value of examining the (usually overlooked) *distribution* of point-wise parameter estimates.

As seen in Figure 6 and as noted above, despite all rats coming from the same breeding batch, the way in which the microstructural parameters changed in response to the training intervention was highly individualized (see Supplementary Figures 2 and 3 for pre- and post-training MTR medians and distributions for each rat). While 9 of the 12 rats showed increased MTR post-training (albeit to different extents), in 3 rats there is an apparent <u>*reduction*</u> in MTR. We hypothesize that such individual differences in changes in myelination measures might reflect individual differences in the adaption of myelin needed to optimize neural synchronies (through fine tuning of conduction velocity) (Pajevic et al., 2014; Chorghay et al., 2018). While axon diameter clearly plays a role in determining conduction velocity (Hursh 1939; Drakesmith et al. 2019), as noted by Pajevic et al. 2014, regulating axon diameter is much less metabolically efficient (and less anatomically plausible over the time-scales considered here) than modulating myelin.

Validating a longitudinal change in myelination is challenging in post-mortem histological data. Only a limited number of rat brains were analyzed by histology (n = 3 pre-training and n = 3 post training). As Table S1 shows (Supplementary Material), there were large inter-individual differences in myelination of white matter tracts at baseline (Coefficient of Variation = 5%), and similar variability in myelination values post-training (Coefficient of variation = 3%); Unlike *in vivo* imaging when multiple longitudinal samples from the same rodent are possible, it is clearly impossible to perform histological assessment of myelin in the same animal pre- and post-training. This, together with the inter-individual variance in both baseline and follow-up myelin measurements reported here, makes it impossible to draw a firm conclusion on the sign and magnitude of any myelin change with such limited histological data. Taking different pairs of pre- and post-training measurements can result in an inference of training-induced increase, reduction or negligible change in myelin. Despite these limitations, we did identify higher average myelin intensities in the post-trained rats in comparison with the pre-trained rats (though not significant when applying a 2 tailed t-test). Future work employing the approach of McKenzie et al. (2014) to block formation of new myelin, in combination with the tractometry approach and myelin-sensitive MRI markers employed here, would seem a more promising way to validate the findings.

In summary, our results demonstrate substantially differential sensitivity of distinct white matter microstructural imaging measures to a spatial working memory training intervention. Taking the median of a given metric along a specific tract neglects a wealth of information, and the *distribution* of parameters may provide a more informative window into white matter plasticity. Moreover, changes in imaging metrics were highly individualized, which may mean that grouping individuals into the same analysis may obscure important changes in microstructure. The most striking differences were seen in the pre-vs-post training distribution of magnetization transfer ratio, specifically in the fornix, which was entirely consistent with our hypothesis concerning the role of the fornix in spatial navigation learning. Overall these results suggest that, while doubtless the most convenient, widely available and most commonly adopted, comparing averaged DT-MRI metrics from within a region of interest is a sub-optimal way to study WM plasticity *in vivo* with the risk of missing important physiological changes.

## Acknowledgements

The data were acquired at the Alfredo Federico Strauss Center for Computational Neuroimaging in Tel Aviv University, funded by a Wellcome Trust Investigator Award (096646/Z/11/Z), which also supported DKJ, DA, CMWT, GDP and DB. BCT was supported by an EPSRC PhD studentship (1367067) and DKJ was supported by a Wellcome Trust Strategic Award (104943/Z/14/Z). We sincerely thank Prof. Mara Cercignani for very useful discussions regarding magnetization transfer analysis and Prof. Richard Bowtell for his support and helpful suggestions regarding quantitative susceptibility mapping.

