## Supplementary Material for "Distributional changes in myelin-specific MRI markers uncover dynamics in the fornix following spatial navigation training"

Bi-lateral fornices, cingulum and corpus callosum were manually reconstructed. Two ROI seeds were chosen on a coronal slice for the fornices reconstruction and another two on the same slice for the cingulum reconstruction. The corpus callosum was reconstructed by choosing an ROI on the mid-sagittal slice.

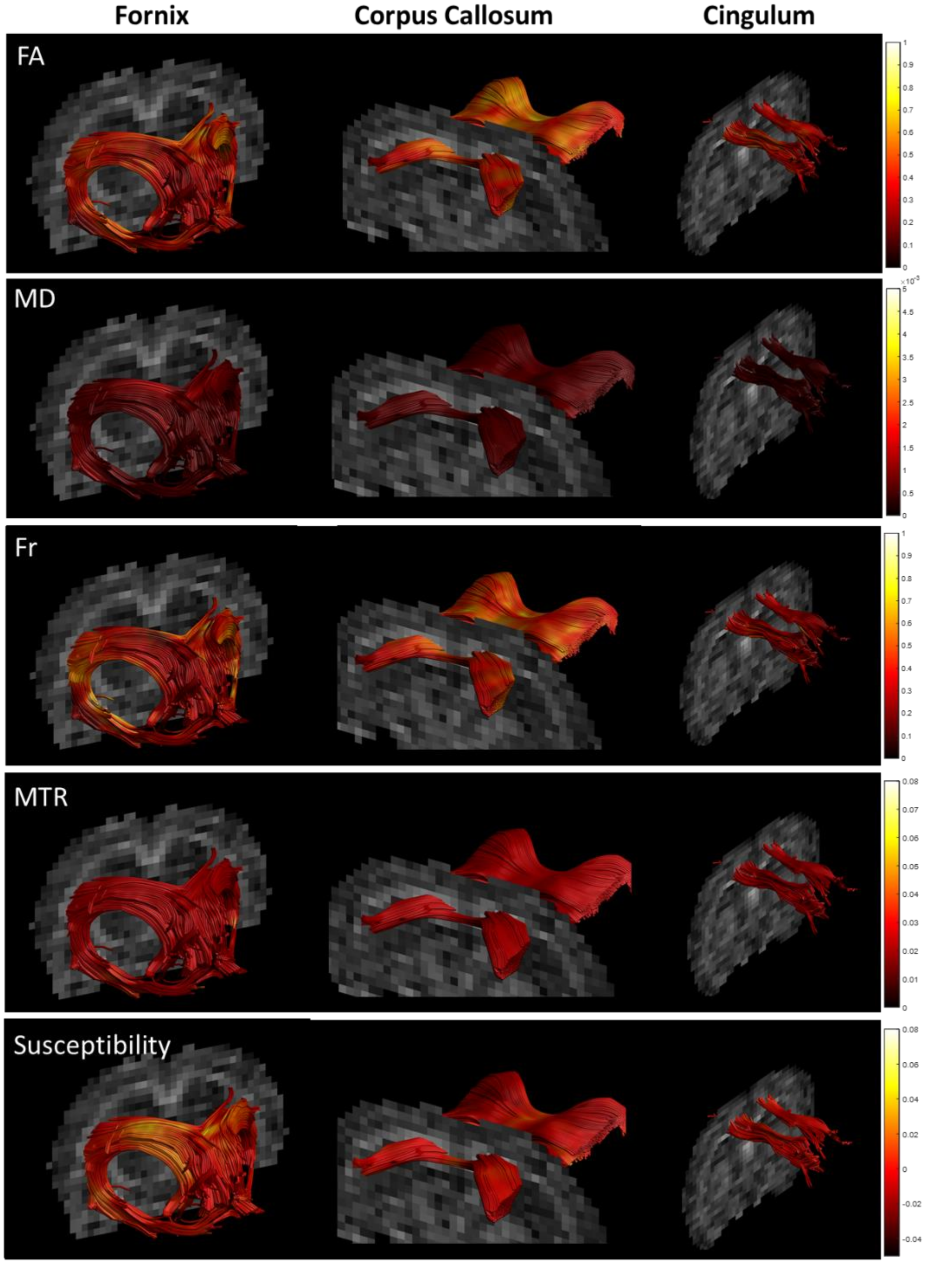


**Figure S1:** Representative parametric maps projected on reconstructions of the three explored tracts; fornix, corpus callosum and cingulum.

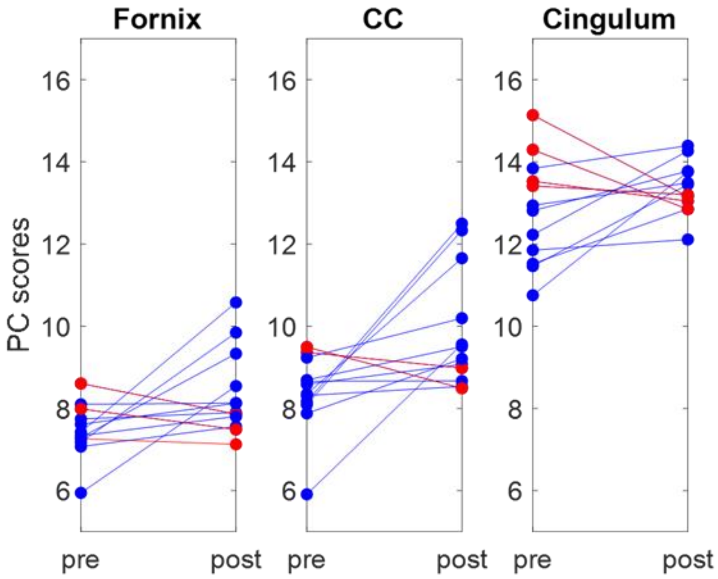


**Figure S2**: Pre- and post-training MTR dominated $PC_5$ scores of individual rats in fornix, CC and cingulum. Blue lines represent scores that increase between pre- and post-training, red lines represent a decrease between pre- and post-training.

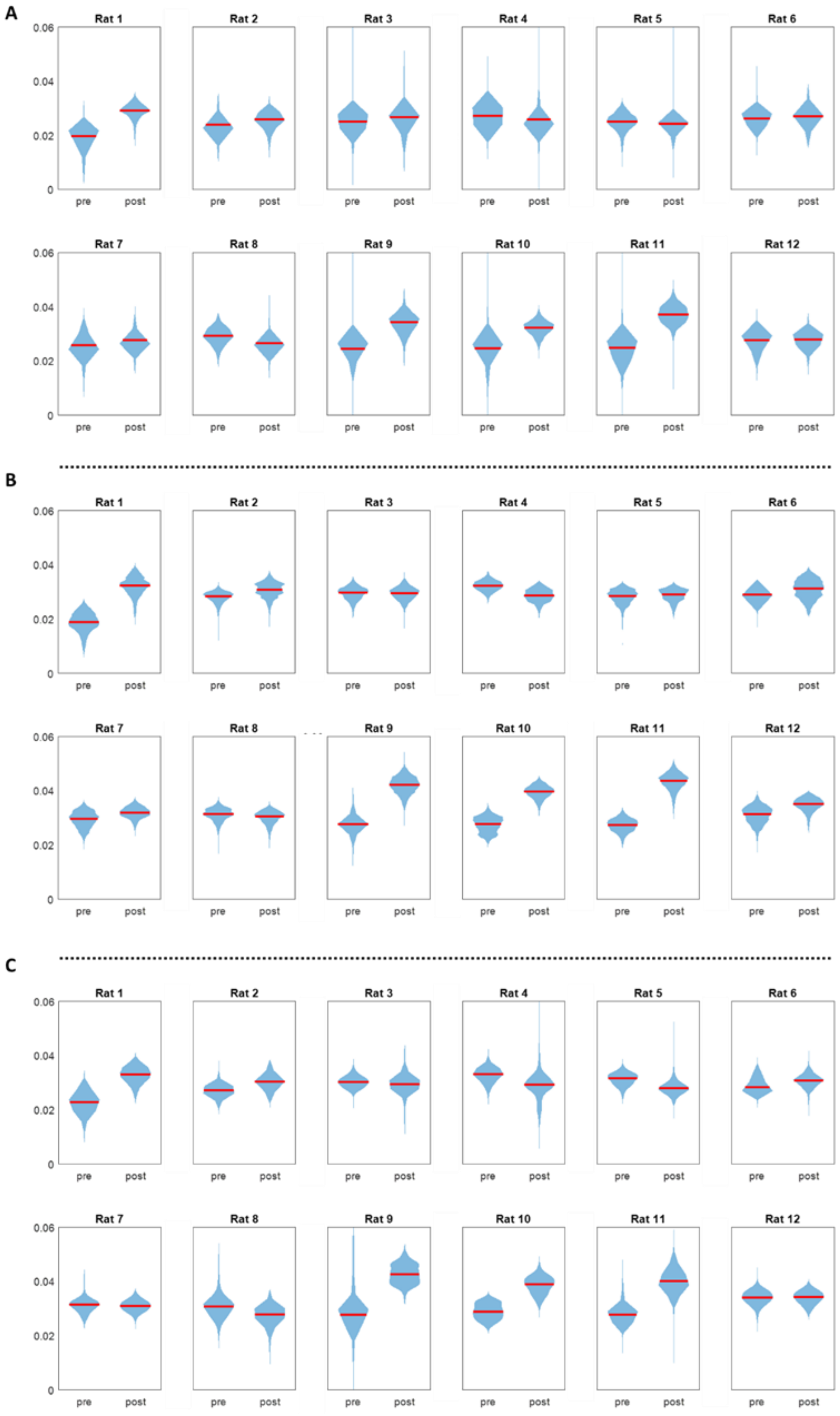


**Figure S3:** Violin plots of MTR values in fornix (A) corpus callosum (B) and cingulum (C), pre- and post-training, for each of the 12 rats. Median is represented by the red lines. Visualizing the distribution of point-wise parameters in this ways provides a more complete picture of individualized responses to training.

**Table S1:** Normalized myelin stain intensity values [a.u.] in six rats (three pre-trained and three post-trained) in the three WM regions of interest. Coefficients of variation [%] are presented in the gray shaded bottom row.

| Fornix | | Corpus callosum | | Cingulum | |
|---|---|---|---|---|---|
| **pre** | **post** | **pre** | **post** | **pre** | **post** |
| 0.52 | 0.54 | 0.52 | 0.52 | 0.52 | 0.53 |
| 0.48 | 0.54 | 0.50 | 0.52 | 0.50 | 0.54 |
| 0.47 | 0.51 | 0.46 | 0.52 | 0.49 | 0.52 |
| **5.12** | **3.16** | **6.66** | **0.41** | **2.58** | **1.87** |
